# Role of Disulfide Bonds and Topological Frustration in the Kinetic Partitioning of Lysozyme Folding Pathways

**DOI:** 10.1101/528000

**Authors:** Aswathy N. Muttathukattil, Prashant Chandra Singh, Govardhan Reddy

## Abstract

Disulfide bonds in proteins can strongly influence the folding pathways by constraining the conformational space. Hen egg white lysozyme has four disulfide bonds and is widely studied for its antibacterial properties. Experiments on lysozyme infer that the protein folds through a fast and a slow pathway. However, the reasons for the kinetic partitioning in the folding pathways are not completely clear. Using a coarse-grained protein model and simulations, we show that two out of the four disulfide bonds, which are present in the *α*-domain of lysozyme, are responsible for the slow folding pathway. In this pathway, a kinetically trapped intermediate state, which is close to the native state is populated. In this state, the orientation of *α*-helices present in the *α*-domain are misaligned relative to each other. The protein in this state has to partially unfold by breaking down the inter-helical contacts between the misaligned helices to fold to the native state. However, the topological constraints due to the two disulfide bonds present in the *α*-domain make the protein less flexible, and it is trapped in this conformation for hundreds of milliseconds. On disabling these disulfide bonds, we find that the kinetically trapped intermediate state and slow folding pathway disappear. Simulations mimicking the folding of protein without disulfide bonds in oxidative conditions show that the native disulfide bonds are formed as the protein folds indicating that folding guides the formation of disulfide bonds. The sequence of formation of the disulfide bonds is Cys64-Cys80 → Cys76-Cys94 → Cys30-Cys115 → Cys6-Cys127. Any disulfide bond, which forms before its precursor in the sequence has to break and follow the sequence for the protein to fold. These results show that lysozyme also serves as a very good model system to probe the role of disulfide bonds and topological frustration in protein folding. The predictions from the simulations can be verified by single molecule fluorescence resonance energy transfer (FRET) or single molecule pulling experiments, which can probe heterogeneity in the folding pathways.

## Introduction

Proteins with disulfide bonds are generally found on the cell surface or extracellular space and they played an important role in elucidating the principles of protein folding.^1–3^ The monumental demonstration that denatured protein chains fold into a unique native functional state under renaturing conditions is established using ribonuclease, which has four disulfide bonds.^4^ A reduced and denatured ribonuclease, which is enzymatically inactive is shown to always fold into a structure with activity equivalent to that of a native protein.^5^ The disulfide bonds in proteins enhance the thermodynamic stability of the folded state and also promote resistance to proteases, which degrades them. The breaking and forming of disulfide bonds are shown to induce conformational changes, which regulate protein activity.^6^

The presence of disulfide bonds can introduce frustration in the folding landscape of evolved proteins, which is funnel-like.^7–9^ Frustration in biomolecular folding can lead to the population of kinetic intermediates resulting in slow folding pathways. ^8^ Proteins inherently have topological frustration, which is due to the requirement that contacts between specific residue pairs should form for the protein to fold while maintaining the chain connectivity. ^8, 10^ Frustration is also observed during the oxidative folding of proteins with disulfide bonds, when a significant part of the protein folds with the premature burial of the thiol group in cysteine before the disulfide bond formation. In these cases, local unfolding of the protein is required for the buried thiol group to get exposed and form the specific disulfide bond present in the protein native state.^2, 3^

In this paper, using coarse-grained protein models and molecular dynamics simulations, we show that the folding of lysozyme with four intact disulfide bonds exhibits kinetic partitioning in its folding pathways^11–13^ due to the frustration introduced by the presence of disulfide bonds. The simulations show that lysozyme folds through a fast and a slow folding pathway in agreement with the experiments.^14–17^ The slow folding pathway is due to the population of a long-lived kinetic intermediate during the late stages of folding. This intermediate survives due to the frustration arising from the restriction in the protein conformational space because of the disulfide bonds, which narrow the folding funnel decreasing the number of escape routes from the trapped state. The slow folding pathway disappears, when lysozyme folding is initiated from an unfolded ensemble without the disulfide bonds, and the formation of disulfide bonds is disallowed mimicking folding in the presence of reducing agents.

Lysozyme due to its antibacterial properties is one of the most widely studied proteins.^19–22^ It is used as a model system to probe the effect of cosolvents^23–30^ and salts^31^ on protein folding and aggregation,^32–34^ study the population of transient intermediates, ^35^ properties of water in the protein hydration shell,^36–38^ and enzymatic activity.^39, 40^ Hen egg white lysozyme is a two domain protein with 129 residues (Figure 1). One domain is rich in *α*-helices and is referred to as the *α*-domain (Lys1 to Asn39 and Ser86 to Leu129), and the second domain contains *β*-sheets and is called the *β*-domain (Thr40 to Ser85). The *α*-domain contains three large *α*-helices (*α*_1_-*α*_3_), each having more than ten residues, and one small helix with six residues (*α*_4_). This domain further includes both the N and C-terminal segments of the protein. The *β*-domain contains triple-stranded anti-parallel *β*-sheets (*β*_1_-*β*_3_) with a long loop. In addition to the *α*-helices and *β*-sheets, there is a short 3^10^ helix in both the domains. The native structure of lysozyme is further stabilized by four disulfide bonds: two in *α* domain (Cys6-Cys127 and Cys30-Cys115), one in *β*-domain (Cys64-Cys80) and another connecting the two domains (Cys76-Cys94).

**Figure 1:**
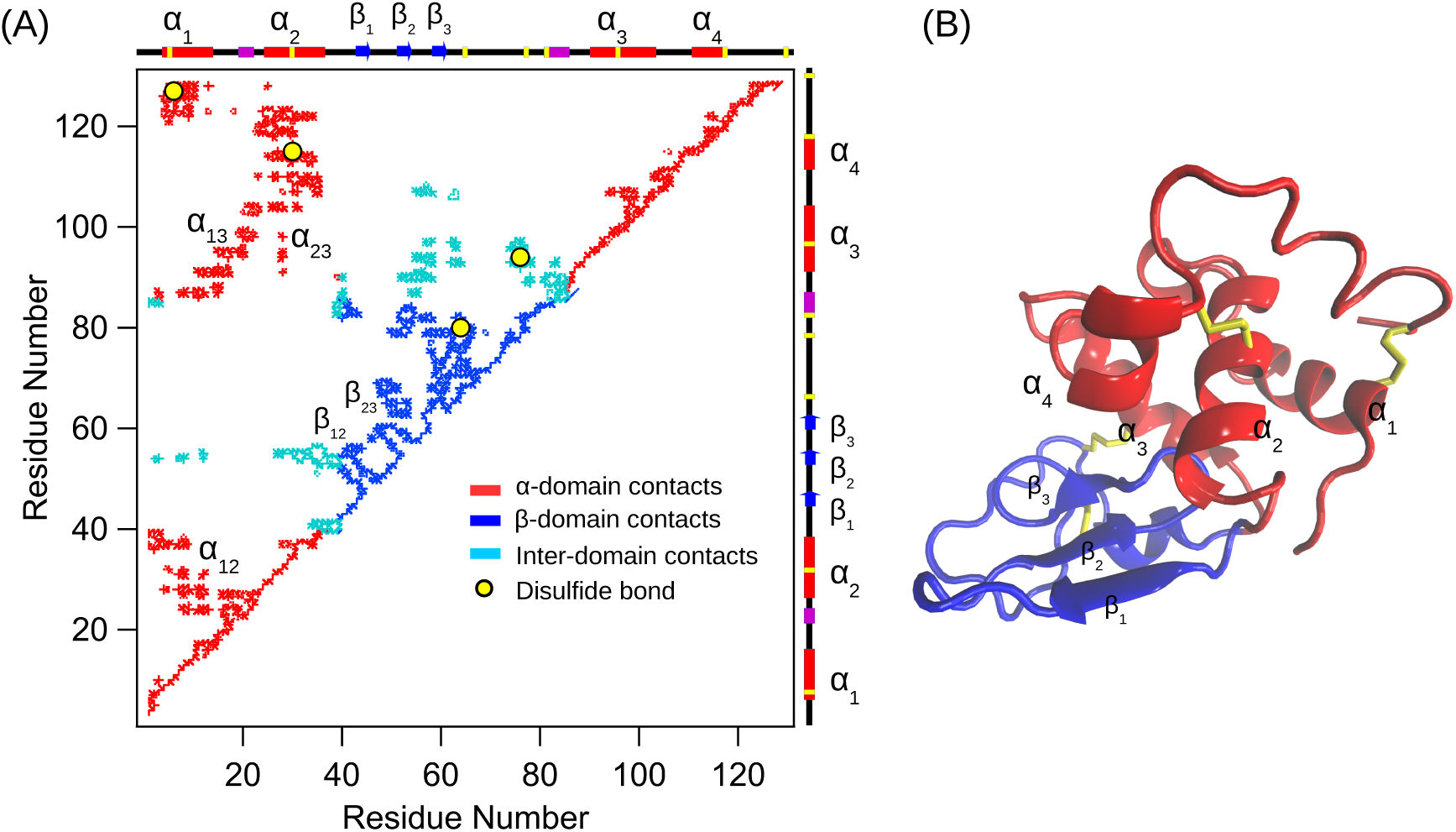
(A) *C_α_* contact map of the native structure of lysozyme. Red, blue and cyan represents contacts between residues present in the *α*-domain, *β*-domain and inter-domain, respectively. The four disulfide bonds are marked as yellow circles. The important secondary and tertiary contacts are marked in the contact map. The secondary structure present along the contour of the protein chain is shown as a schematic along the right and upper axes: *α*-helices are red boxes, 3^10^-helices are purple boxes and *β*-strands are blue arrows. (B) Cartoon representation of the crystal structure^18^ of lysozyme (protein data bank (PDB) ID: 1AKI). *β*-domain, *α*-domain and disulfide bonds are shown in blue, red and yellow, respectively.

A number of experiments have inferred that lysozyme with four intact disulfide bonds folds through a fast and a slow folding pathway.^14–17^ These experiments have also concluded that an intermediate is populated in the slow folding pathway.^41^ However, a complete understanding of the slow folding pathway is still lacking. The reasons for kinetic partitioning in the folding pathways are unknown and the structure of the intermediate populated in the slow folding pathway is not available. Questions such as how do the disulfide bonds influence lysozyme folding pathways, and whether protein folding guides disulfide formation or vice versa during oxidative folding still persist.^42–44^ To address these questions we performed Brownian dynamics simulations to study lysozyme folding kinetics using the coarse-grained self-organized polymer-side chain (SOP-SC) model.^45, 46^ To comprehensively understand the role of disulfide bonds in lysozyme folding, we performed simulations mimicking three different conditions. In the first set of simulations, we studied lysozyme folding with all the four intact disulfide bonds, in the second set we disabled the disulfide bond formation mimicking lysozyme folding in reduced conditions, and in the third set the disulfide bonds form as the protein folds mimicking oxidative folding.

## Results and Discussion

We spawned 100 Brownian dynamics simulation trajectories to probe the folding of lysozyme with four intact disulfide bonds at *T* = 320 K (*≈* 0.8 *T_M_*, where *T_M_* is the melting temperature). A detailed description of the coarse-grained protein model and simulation methods are in the supporting information (SI). In these conditions, the radius of gyration 〈*R_g_*〉 of the protein as a function of time for different trajectories shows that lysozyme folds through a fast and a slow folding pathways exhibiting kinetic partitioning^12–17^ (Figure 2). In 49 out of the 100 folding trajectories, lysozyme folds through a fast folding pathway (Figure 2A). In rest of the trajectories, lysozyme folds through a slow folding pathway, where a partially folded intermediate state is populated (Figure 2B). The average folding time of the fast folding trajectories is 9.7 ± 4.0 ms, and in contrast, the folding time for the slow trajectories is on the order of hundreds of milliseconds, which is in agreement with the experiments.^15, 47^ In 49 out of the 51 slow folding pathways, the protein did not fold completely on a time scale of 500 ms, which is the maximum run time of the simulations, and is trapped in a conformation with *R_g_ ∼* 17 Å for hundreds of milliseconds (Figure 2B).

**Figure 2:**
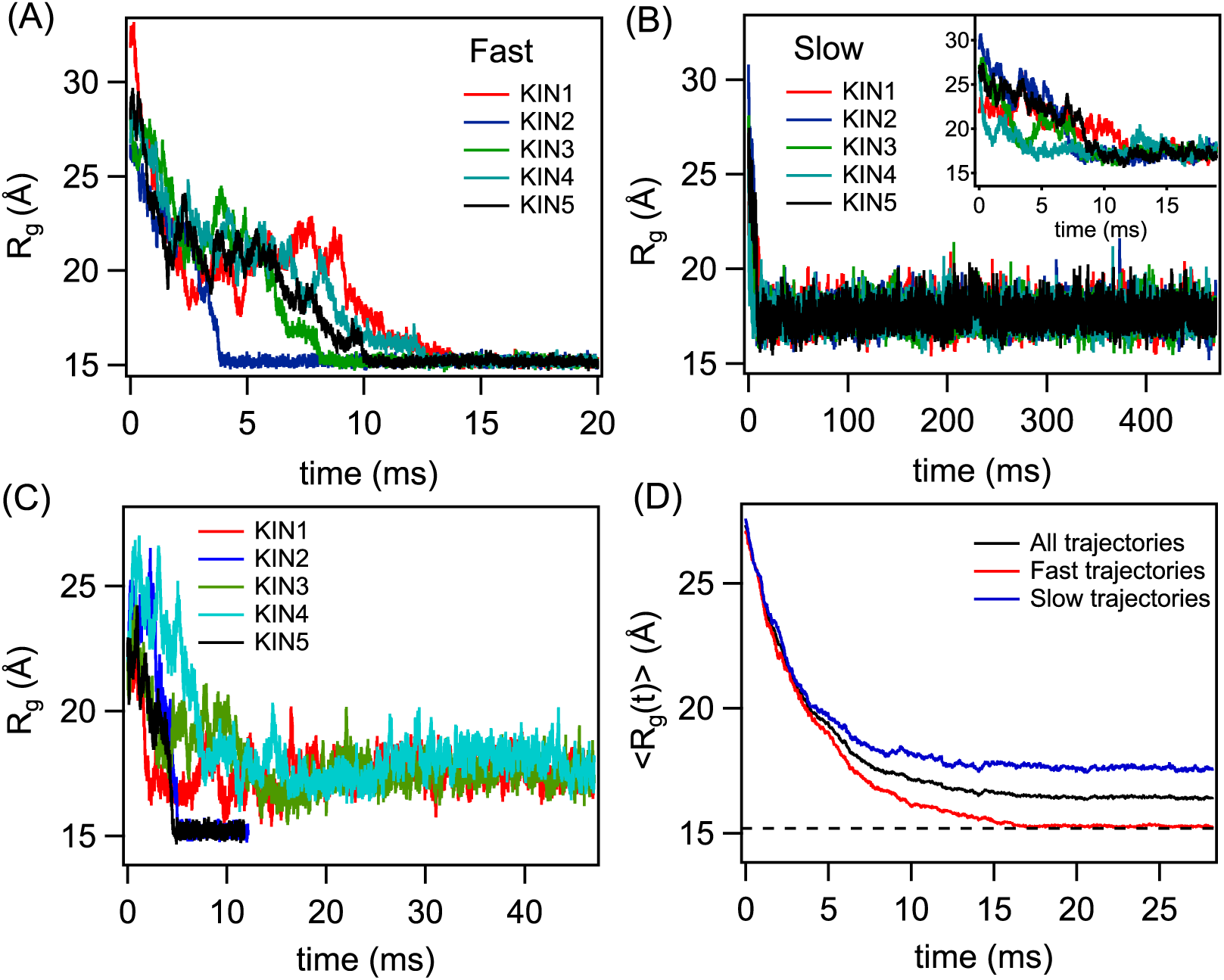
Representative simulation trajectories of fast (A) and slow (B) folding pathways of lysozyme. Time evolution of the radius of gyration (*R_g_*) shows that in fast folding pathways, protein folds to the native state (*R_g_* ≈ 15 Å) within 15 ms. In the slow pathways, protein does not fold within 500 ms of the simulation time. In the slow pathways, protein is found to be kinetically trapped in a state with *R_g_* ≈ 17 Å. The initial stages of folding in slow trajectories are shown in the inset of panel-B. (C) *R_g_* plotted as a function of time for the five Brownian dynamics folding simulations spawned using the same initial unfolded conformation. Trajectories KIN2 and KIN5 fold within 10 ms using the fast folding pathway. The remaining three trajectories, KIN1, KIN3, and KIN4, fold through the slow folding pathway and are trapped in the intermediate state. This shows that stochasticity in the folding dynamics is responsible for kinetic partitioning in lysozyme folding pathways. (D) Time evolution of the ensemble average of *R_g_*, (〈*R_g_*(*t*)〉) of all the folding trajectories. 〈*R_g_*〉 as a function of time for the subset, fast and slow trajectories, is also shown. The dotted line shows *R_g_* of the protein in the native state. All the fast trajectories (49 out of 100) fold to the native state within ∼ 15 ms, while slow trajectories (51 out of 100) are trapped in a conformation with *R_g_ ∼* 17 Å.

The initial unfolded conformation in all the folding trajectories are different, and these conformations are generated using a simulation performed at temperature, *T* = 420 K (> *T_M_*) (Figure S1). To rule out the possibility that the kinetic partitioning is due to a heterogeneous mixture of unfolded molecules, ^15, 48^ *i.e.* one set of unfolded molecules fold through the fast pathway and a second set folds through the slow pathway, we also performed multiple folding simulations starting from the same unfolded conformation. Even in this case, the protein folds using both the slow and fast folding routes exhibiting kinetic partitioning (Figure 2C). Thus, it is the stochasticity in the folding dynamics, which is responsible for the protein choosing either the fast or slow folding route. The simulations broadly support the triangular folding mechanism proposed by Guo and Thirumalai,^13^ and Wildegger and Kiefhaber,^16^ with a trapped kinetic state as shown below

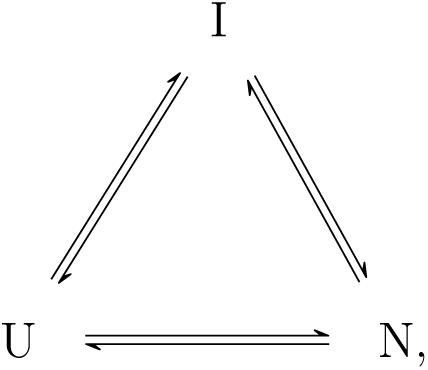

where N, I and U refer to lysozyme native, intermediate and unfolded states, respectively.

On initiating folding, lysozyme forms compact structures in both the fast and slow folding pathways within ∼ 5 ms (Figure 2D). During this compaction, the average radius of gyration, 〈*R_g_*〉, of lysozyme decreases by *≈* 8 Å. The compact state has an average *R_g_* of ∼ 20 Å, which is in agreement with the experimentally^49^ measured value of 19.6 Å. The formation of *β*-domain (Figure 3A) and helices (Figure 3B, S2) contribute to the compaction in protein dimensions. The compacted state mainly lacks the tertiary contacts present in *α*-domain and the inter-domain contacts (Figure S3). After the compaction, bifurcation in the folding pathways to fast and slow folding routes is observed. In the fast folding pathway, the protein continues to fold to its native structure with *R_g_ ≈* 15 Å, on an average time scale of ∼ 15 ms with the formation of inter-domain contacts. ^49^ However, in the slow folding pathways, the protein gets trapped in an intermediate state with lifetimes on the order of hundreds of milliseconds. The *R_g_* of this intermediate state is *≈* 17 Å, and it is in good agreement with the experimentally^49^ measured value of 16.7 Å. The *R_g_* value of the intermediate state indicates that the protein is close to the folded state (Figure 2D).

**Figure 3:**
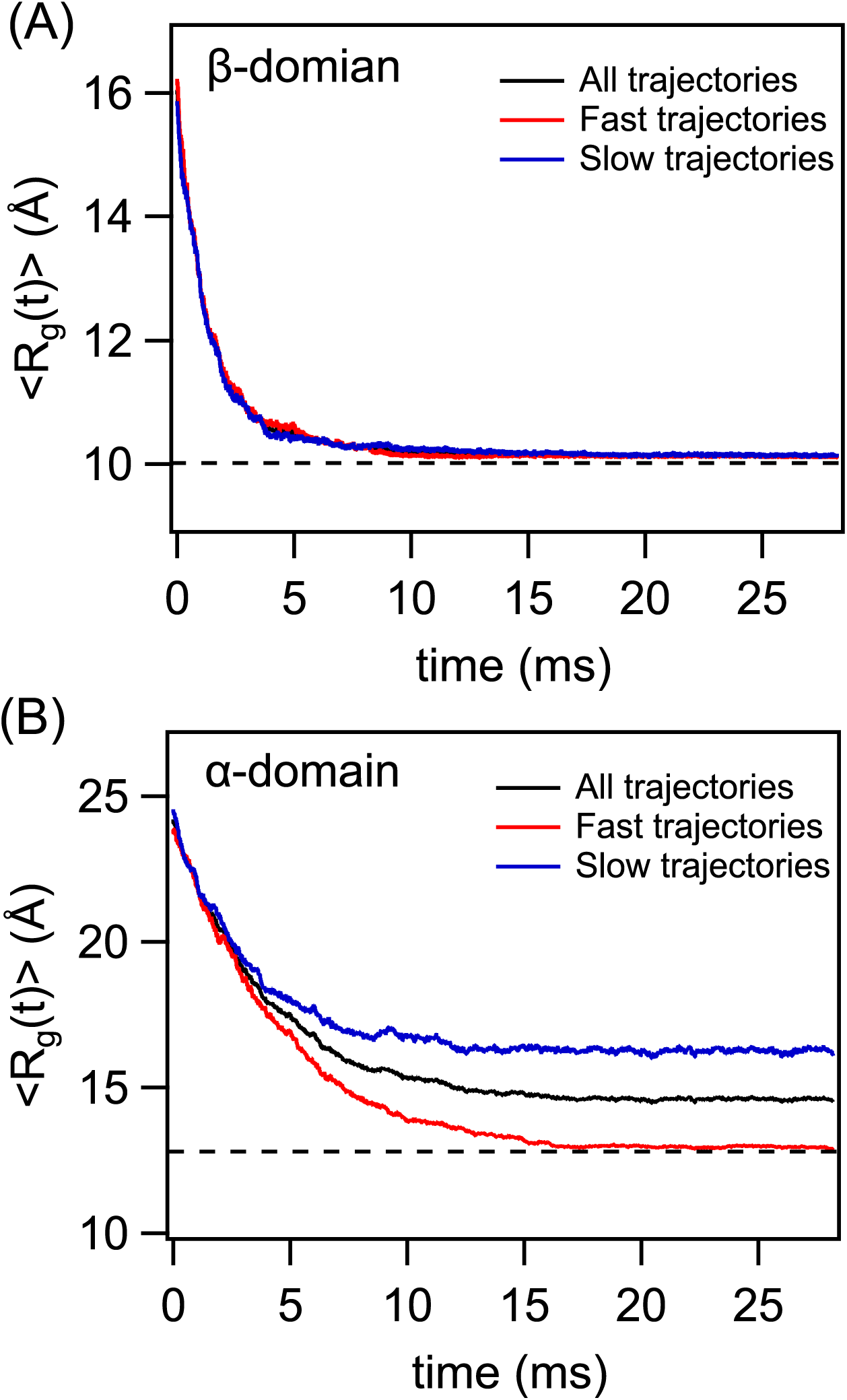
Time evolution of the ensemble average of radius of gyration (〈*R_g_*(*t*)〉) of *β*-domain (A) and *α*-domain (B) in fast and slow folding trajectories. *β*-domain folds fast in both the pathways, while *α*-domain shows differences in the folding kinetics between fast and slow pathways. The dotted line shows *R_g_* of the respective domains in the native state.

To probe the folding of *α* and *β*-domains, we computed 〈*R_g_*〉 of these individual domains as a function of time. In both the fast and slow folding pathways, the *β*-domain folds completely within 5 ms reaching the native state 〈*R_g_*〉 value of ∼ 10 Å (Figure 3A). The time evolution of 〈*R_g_*〉 in the folding of *α*-domain shows that it is the rate determining step in the folding of lysozyme. In the fast folding trajectory, the *α*-domain folds on a time scale of ∼ 15 ms reaching the native state 〈*R_g_*(*t*)〉 value of ∼ 12.5 Å (Figure 3B). Whereas in the slow folding trajectories, the *α*-domain is stuck in an intermediate state with a 〈*R_g_*〉 value of ∼ 17.5 Å (Figure 3B). This shows that *α*-domain folding in lysozyme is responsible for the population of slow folding pathways in lysozyme folding.

In the folding of *α*-domain, the three large *α*-helices (*α*_1_-*α*_3_) influence the folding kinetics. In both the fast and slow folding pathways, the formation of individual helices precedes the formation of inter-helical contacts. In 70% of the fast folding pathways, we find that the inter helical contacts between *α*_2_ and *α*_3_ (*α*_2_*α*_3_) form first, and then the inter helical contacts *α*_1_*α*_2_ and *α*_1_*α*_3_ form almost simultaneously along with the inter-domain contacts between the *α* and *β*-domains leading to the complete folding of the protein (Figure 4A). In the remaining 30% of fast folding trajectories, *α*_1_*α*_2_ contacts form first, followed by the simultaneous formation of *α*_2_*α*_3_, *α*_1_*α*_3_ and inter domain contacts (Figure 4B). In the fast folding trajectories, either *α*_2_*α*_3_ or *α*_1_*α*_2_ contacts form first followed by the simultaneous formation of other two sets of inter helical contacts and inter domain contacts. This mechanism of formation of interhelical contacts with the orientation of the helices similar to the orientation present in the native state makes lysozyme folding smooth without any kinetic traps.

**Figure 4:**
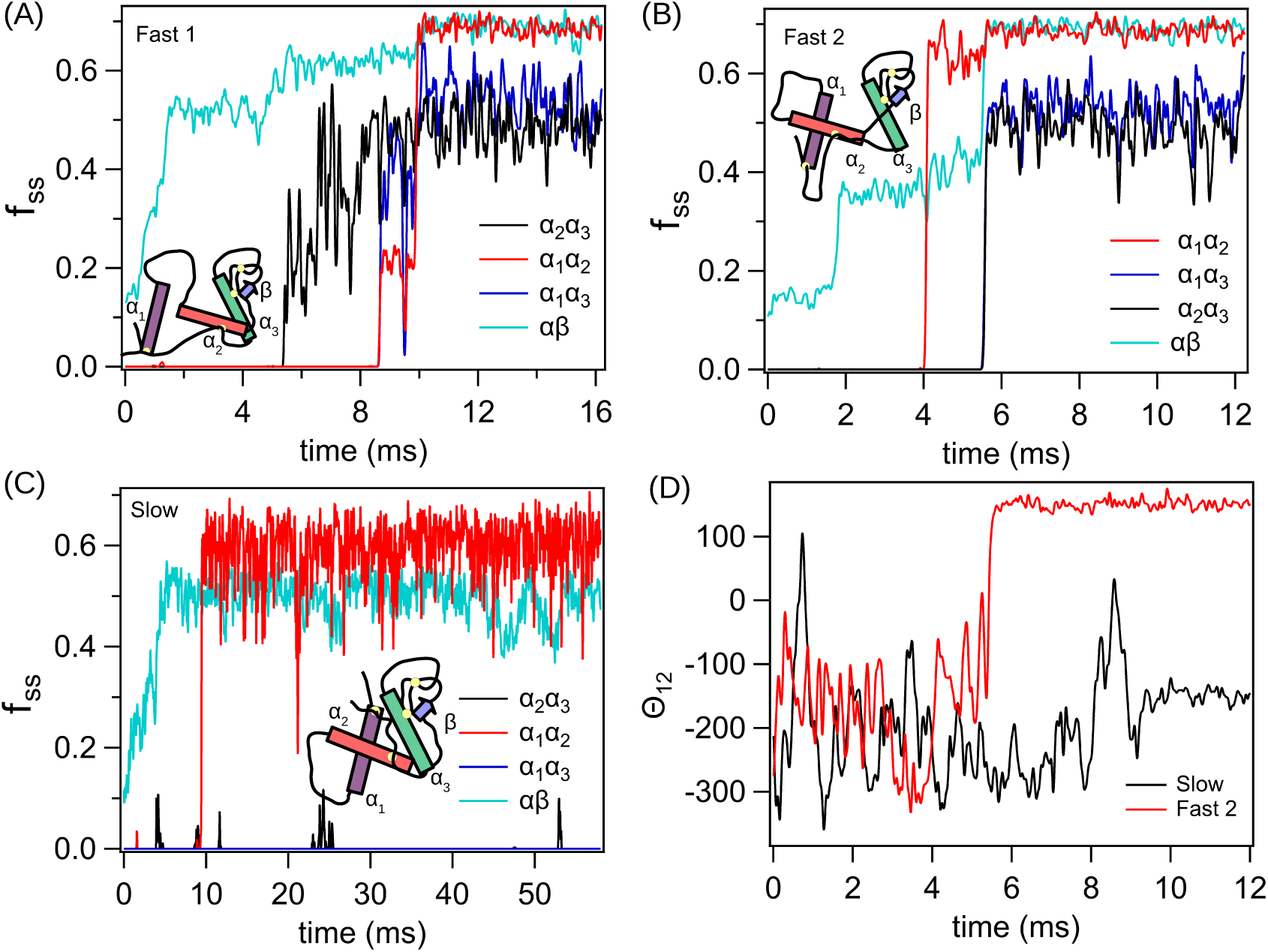
Time evolution of the fraction of native inter-helical (*f_αα_′*) and inter-domain (*f_αβ_*) contacts in representative fast (A and B) and slow (C) folding trajectories. (A) In 70% of the fast folding trajectories, *α*_2_*α*_3_ contacts form first followed by the formation of *α*_1_*α*_2_, *α*_1_*α*_3_ and *αβ*. (B) In the remaining 30% of the fast folding trajectories, *α*_1_*α*_2_ contacts form first followed by the formation of *α*_1_*α*_3_, *α*_2_*α*_3_ and *αβ*. (C) In the slow folding pathways, the protein is trapped in a kinetic intermediate, and in this state a significant fraction of *α*_1_*α*_2_ and *αβ* contacts are formed. (D) The relative orientation of *α*_1_ and *α*_2_ helices in fast (panel-B) and slow (panel-C) folding pathways is shown using the scalar triple product, 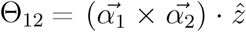, where 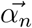 is the vector along *α_n_* and 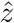 is the unit vector along *z*-axis. In both, fast and slow folding pathways (panel-B and C), *α*_1_*α*_2_ contacts form early, but their relative orientations are different. In the fast pathways, the relative orientation of *α*_1_ and *α*_2_ is similar to that of native state.

In the slow folding pathways, a significant fraction of *α*_1_*α*_2_ contacts form first (Figure 4C). This structure also prevents the formation of *α*_2_*α*_3_ and *α*_1_*α*_3_ contacts. By visually observing this trapped conformation, we noticed that the orientation of helix *α*_1_ is in the opposite direction relative to the helices *α*_2_ and *α*_3_ compared to the native folded state (see the schematics in the inset of Figures 4B, C). For the protein trapped in this state to fold correctly, *α*_1_ has to reorient by breaking the *α*_1_*α*_2_ contacts, and this leads to a long-lived kinetic intermediate (Figure 4D, S2). To quantitatively show that *α*_1_ is misaligned relative to *α*_2_, we computed the scalar triple product 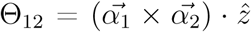, where 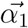 is the vector joining the backbone beads of residues Arg5 and Arg14 in 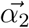 is the vector joining the backbone beads of the residues Leu25 and Ser36 in *α*_2_, and 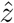 represents the unit vector along *z*-axis. The protein conformations before computing Θ_12_ are aligned to the native state. The plot of Θ_12_ as a function of time for the fast and slow folding pathways clearly shows that the helices *α*_1_ and *α*_2_ have a different orientation. The contact map of kinetically trapped intermediate state shows that the *β*-domain is completely folded, and *α*-domain is partially folded with *α*_2_*α*_3_, *α*_1_*α*_3_ and some of the inter-domain contacts missing (Figure 5). The population of a misfolded intermediate leading to a slow lysozyme folding pathway was also speculated using a model four helix bundle by Thirumalai and Guo.^12^

**Figure 5:**
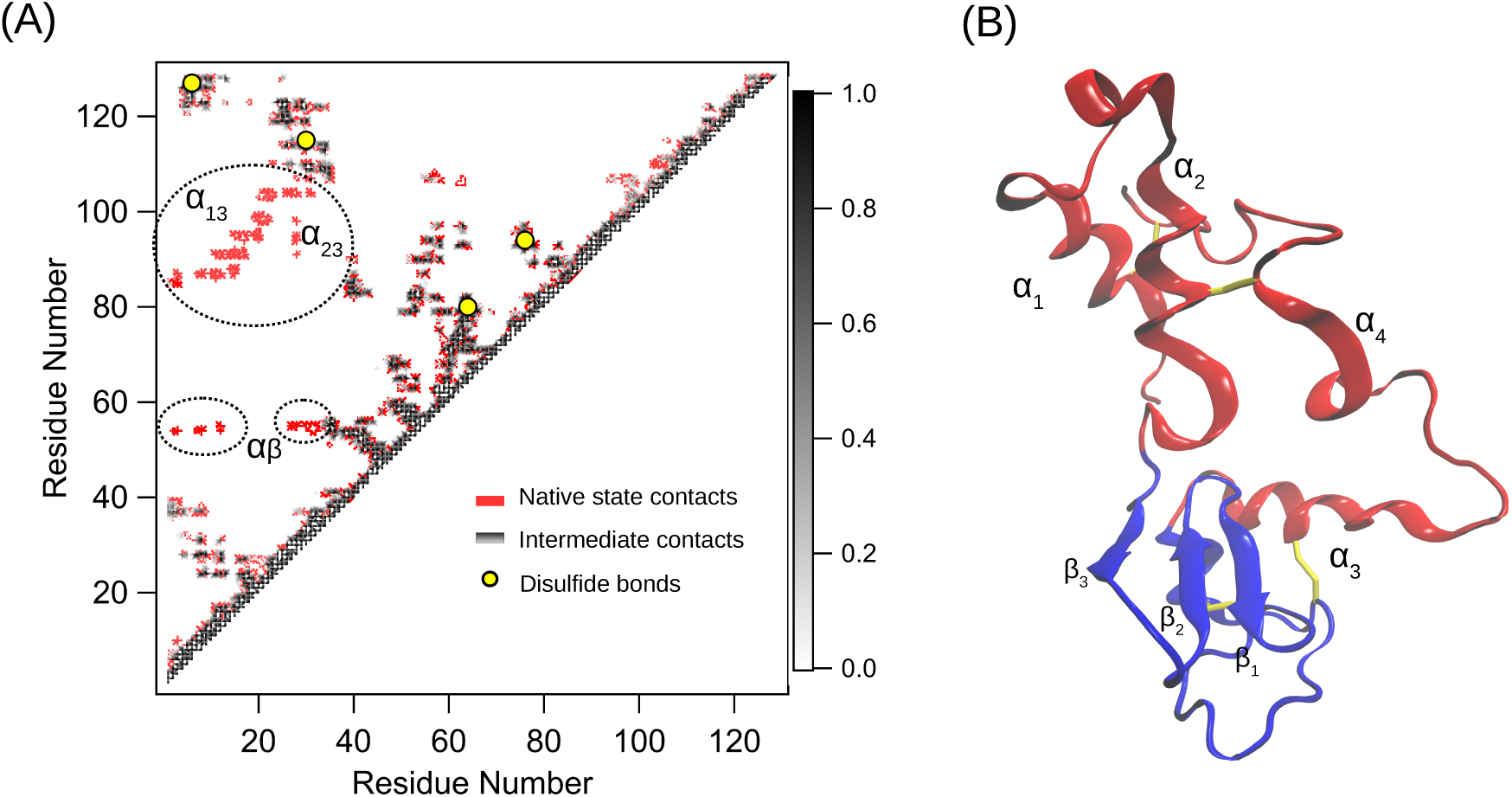
(A) *C_α_* contact map of the kinetic intermediate populated in the slow folding pathways. Contacts between the residues present in the native and intermediate state are shown in red and gray. The inter helical contacts, *α*_1_*α*_3_ and *α*_2_*α*_3_, and some of the inter-domain contacts (*αβ*) are missing in the intermediate state compared to the native state. (B) Representative structure of the kinetically trapped intermediate in the slow folding trajectories. The *α* and *β*-domains are shown in red and blue, respectively. In the intermediate state, *β*-domain is folded and *α*-domain is partially folded.

At 320 K, we observed complete folding of lysozyme in 2 out of the 51 slow folding trajectories within the simulation time of 500 ms. The simulations show that the kinetically trapped misfolded intermediate has to partially unfold before folding back to the native state as inferred in some of the experiments^50, 51^ (Figure 6, S4). In these trajectories, the *α*_1_*α*_2_ contacts present between the misaligned helices in the intermediate state break down, allowing the protein to escape from this trapped state (Figure 6A). The disruption in the *α*_1_*α*_2_ contacts, allows *α*_1_ to reorient to the correct alignment relative to *α*_2_, and the protein folds back to the native state (Figure 6B). If disruption in the *α*_1_*α*_2_ contacts is a primary bottle neck for the protein to escape from this trapped state, then increased thermal fluctuations at a higher temperature should facilitate the protein to escape from the trapped state. To verify this we performed additional 50 Brownian dynamics simulations of lysozyme folding at a higher temperature, *T* =380 K (< *T_M_*). Out of the 50 trajectories, the kinetically trapped intermediate state is populated in 14 trajectories. In 10 out of the 14 trajectories, on a time scale of 200 ms, the protein transitioned from the intermediate state by disrupting the *α*_1_*α*_2_ contacts, as observed in the simulations performed at the lower temperature (Figure 6, S4). Even in experiments,^50^ the folding of lysozyme at a higher temperature is found to be more cooperative because the thermal fluctuations at a higher temperature will facilitate the protein to easily unfold locally and escape from the kinetic intermediate state. The population of the intermediate state is not due to the formation of non-native interactions as hypothesized,^51, 52^ as the coarse-grained native-centric protein model^45, 46, 53^ used in this study has no energetic frustration. In this model, non-native interactions cannot form, as all the pairs of residues, which are not native, interact only with a repulsive potential (see SI for more details). As a result, only topological frustration, and no energetic frustration, exists in these models. This shows that it should be the topological frustration and with a possible role of the disulfide bonds, which constrain the conformational space available for the protein should facilitate the population of the kinetically trapped intermediate state.

**Figure 6:**
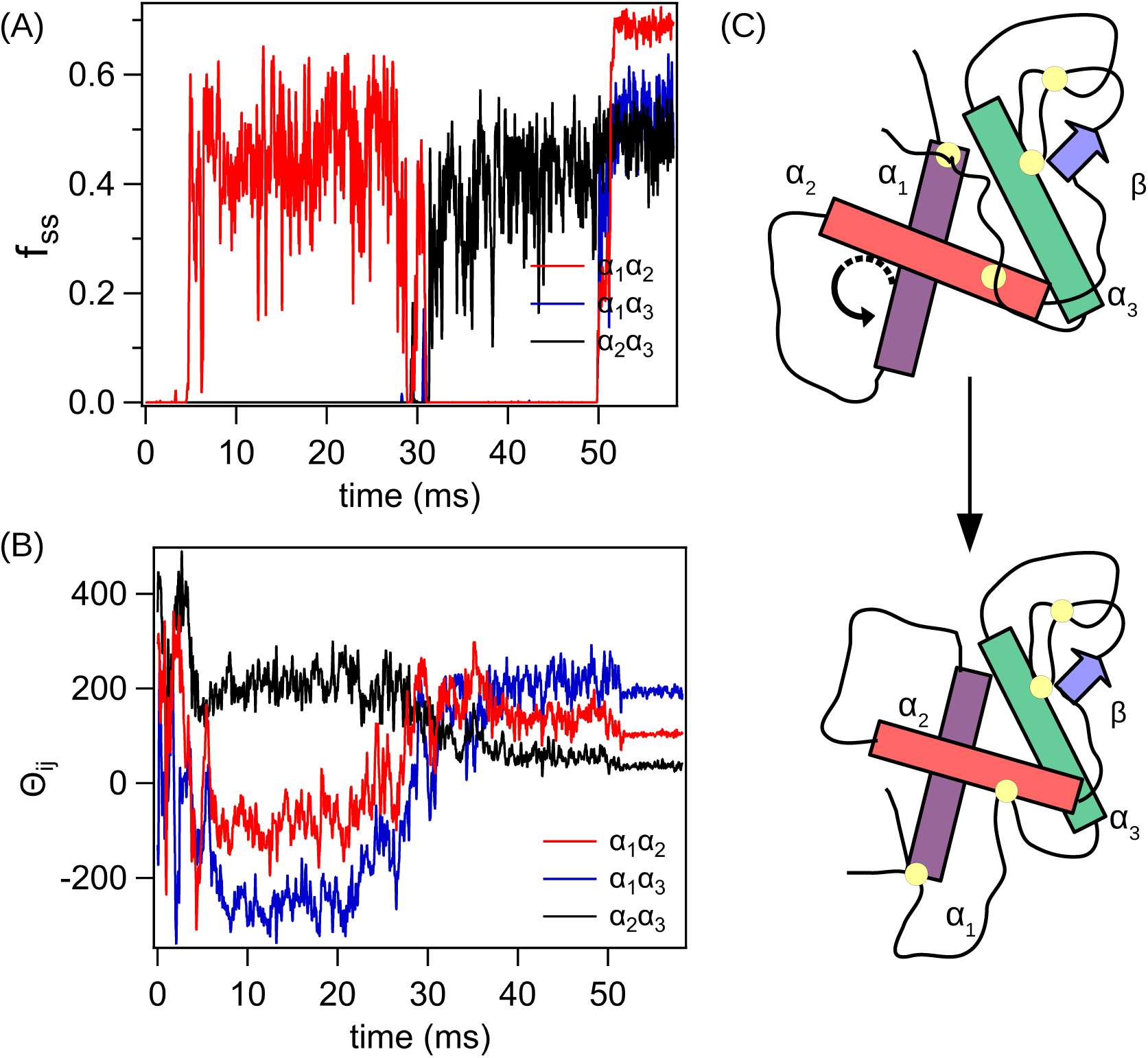
(A) A representative slow folding trajectory, where the protein escapes from the kinetically trapped state and folds to the native state. The *α*_1_*α*_2_ contacts between the helices in the incorrect orientation fall apart at 30 ms, allowing the protein to fold back to the native state. (B) The relative orientation of helices quantified using the scalar triple product, 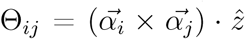, where 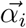 is the vector along *α_i_* and 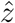 is the unit vector along *z*-axis, shows that *α*_1_ is incorrectly oriented relative to *α*_2_ and *α*_3_ in the kinetically trapped state. As the *α*_1_*α*_2_ contacts fall apart at 30 ms, the orientation changes and the protein folds to the native state. (C) Schematic showing the orientation of the helices in the trapped and native state. *α*_1_ orientation in trapped and native states is different relative to *α*_2_ and *α*_3_. The residues forming the disulfide bonds are shown as yellow dots.

To test the possible role of disulfide bonds in the population of kinetic intermediate, we performed additional 100 folding simulations of lysozyme without the presence of disulfide bonds at the same temperature *T* = 320 K. This mimics the folding of lysozyme in the presence of reducing agents such as dithiothreitol, which prevents the formation of disulfide bonds. In these simulations we did not observe any kinetically trapped intermediate states and all the trajectories reached the folded state within 200 ms (Figure 7A). This shows that in lysozyme conformations, where *α*_1_ is misaligned relative to *α*_2_ (Figure 5), the topological constraints due to the disulfide bonds makes the protein less flexible for structural interconversions, and these conformations result in kinetic traps in the folding pathways. To escape from this conformation, local unfolding of lysozyme, which is the unwinding of *α*_1_*α*_2_ inter-helical contacts is required. In renaturing conditions, the time scales to cross the barrier for local unfolding is on the order of hundreds of milliseconds, and this results in the slow folding pathway of lysozyme. The population of a misfolded kinetic intermediate due to topological frustration, which requires local unfolding or back tracking to escape from the kinetic trap is also observed in the folding of interleukin-1*β*,^54, 55^ green fluorescent protein (GFP),^56, 57^ cytochrome *c*^58^ and a model four helix bundle.^12, 13^

**Figure 7:**
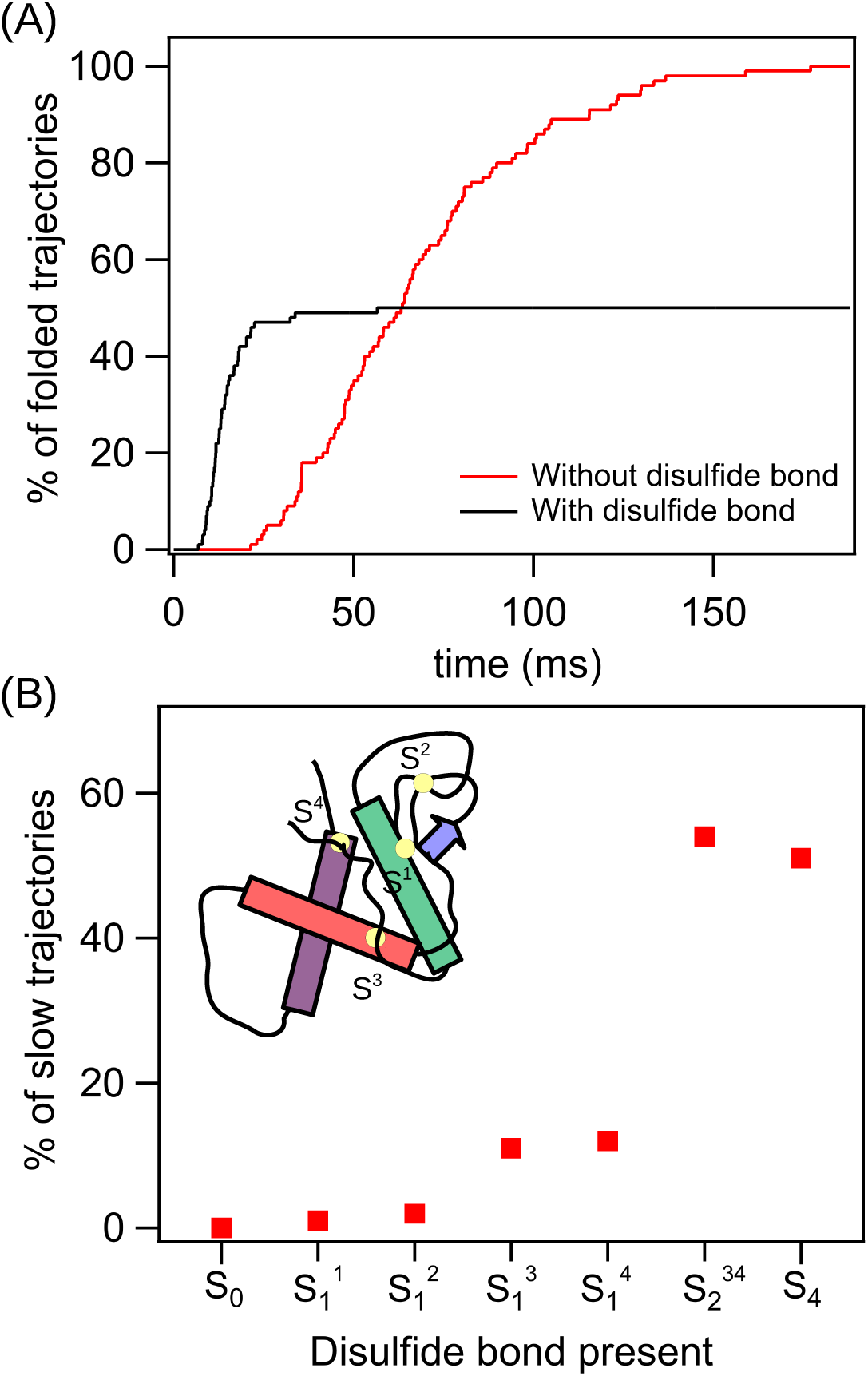
(A) Percentage of simulation trajectories, where the protein has folded as a function of simulation time. All the simulation trajectories fold within 200 ms when the disulfide bonds are absent, and their formation is disabled. In the presence of all the four disulfide bonds, ~50% of the trajectories fold, and the rest are stuck in a kinetically trapped intermediate state. This shows that disulfide bonds are responsible for the slow folding pathway and population of the kinetic intermediate. (B) Percentage of slow folding trajectories where kinetically trapped intermediate state is populated when different combinations of disulfide bonds are allowed in the protein. S_0_ and S_4_ represent the protein with zero disulfide bonds, and all the four disulfide bonds present, respectively. 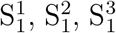 and 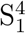 represent the protein with only one disulfide bond Cys64-Cys80, Cys76-Cys94, Cys6-Cys127, and Cys30-Cys115, respectively. Slow folding pathways are observed in proteins with Cys6-Cys127 or Cys30-Cys115 disulfide bonds. The number of slow trajectories observed when these two disulfide bonds (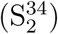) are present is equal to the number of slow trajectories observed when all the four disulfide bonds are present. This implies the disulfide bonds Cys6-Cys127 and Cys30-Cys115 present in the *α*-domain are responsible for the population of the kinetically trapped intermediate resulting in slow folding pathway.

To probe the role of individual disulfide bonds in the population of the intermediate state, we performed four sets of simulations and in each set only one particular disulfide bond is kept intact and the other three bonds are disabled. In each set, we spawned 50 folding trajectories at *T* = 320 K. The simulations clearly shows that the two disulfide bonds, Cys6-Cys127 and Cys30-Cys115, present in the *α*-helical domain of the protein are responsible for the kinetic partitioning in the folding pathways and facilitate the population of the kinetic intermediate in the slow folding pathway in agreement with the experiments^59–61^ (Figure 7B). When only these two disulfide bonds are present, the system shows a similar percentage of the population of slow trajectories compared to that of a protein with all the four disulfide bonds present. This observation is completely in agreement with the experiments of Giez *et al.*,^61^ and since the population of the kinetic intermediate is due to the restriction in the rearrangement of helices *α*_1_ and *α*_2_, and the two disulfide bonds, Cys6-Cys127 and Cys30-Cys115, are attached to these helices.

Studies^42, 43^ show evidence that protein folding guides disulfide bond formation. We performed simulations mimicking oxidative folding of lysozyme to probe the sequence of disulfide bond formation, and whether protein folding guides disulfide bond formation or vice versa. We used the method developed by Qin *et al.*^43^ to mimic the disulfide bond formation. In this method, we assume that a disulfide bond is formed when three criteria related to proximity, orientation and solvent exposure of cystine residues are satisfied (see SI for details). When this criteria is satisfied, a FENE potential and an angle potential corresponding to the disulfide bond is added to the side chains of the cystine residues mimicking the bond formation.

We have spawned 100 lysozyme folding trajectories at *T* = 320 K mimicking folding in oxidative conditions. In all the folding trajectories, we find that Cys64-Cys80 disulfide bond present in the *β*-domain forms within 10 ms after folding is initiated (Figure 8). The second disulphide bond formed is the Cys76-Cys94, and it is located at the interface of the *α* and *β*-domains. The disulfide bonds, Cys64-Cys80 and Cys76-Cys94, are short-ranged native contacts between the side chains of the Cys residues forming these bonds. The number of residues separating the Cys residues forming the bonds along the contour of lysozyme chain is less than 18. As a result, during the initial protein collapse, these Cys residues come closer and become solvent inaccessible leading to the formation of the disulfide bond.

**Figure 8:**
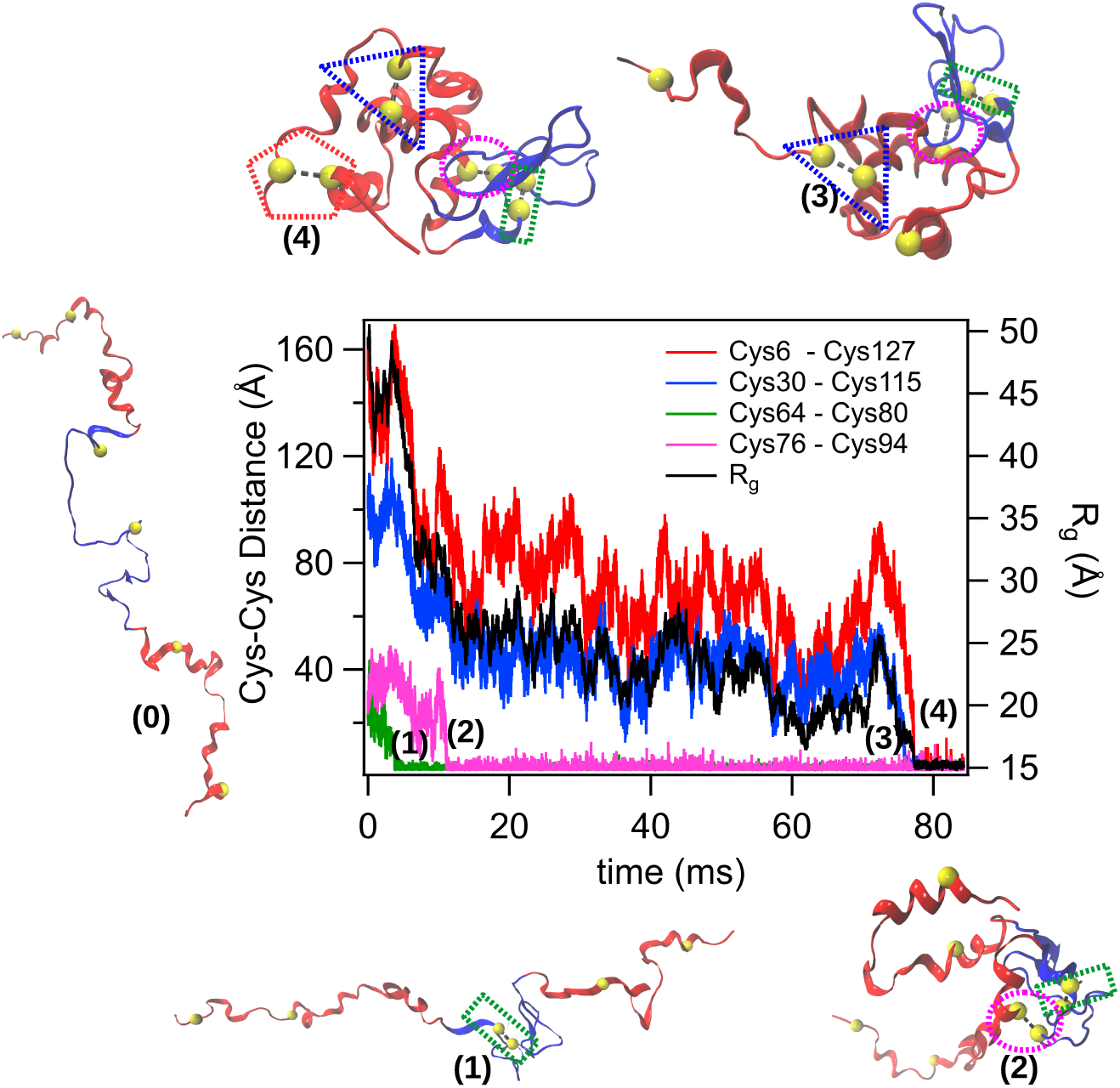
A representative folding trajectory of lysozyme in oxidative folding conditions. In the initial unfolded conformation, none of the disulfide bonds are present. The distance between various pairs of Cys residues, which form disulfide bonds in the native state are plotted as a function of time. The sequence of disulfide bond formation is as follows: Cys64-Cys80 → Cys76-Cys94 → Cys30-Cys115 → Cys6-Cys127. Representative snapshots of lysozyme conformations show the formation of disulfide bonds as the protein folds.

The other two disulfide bonds, Cys30-Cys115 and Cys6-Cys127, which form in the last stages of folding are in the *α*-domain, and these bonds are long-ranged native contacts between the side chains of the Cys residues forming these bonds, as they are well separated along the contour of the lysozyme chain. The Cys30-Cys115 and Cys6-Cys127 disulfide bonds are either formed simultaneously (only 6 % - Figure S4), or Cys30-Cys115 is formed before Cys6-Cys127. The order of disulfide bond formation in lysozyme during oxidative folding is Cys64-Cys80, Cys76-Cys94, Cys30-Cys115, and Cys6-Cys127.

If any one of the disulfide bonds form before its precursor in the order, then the protein does not fold to its native state (Figure S5). The disulfide bond, which has formed prematurely in the order gets solvent exposed, *i.e.* in this case, the bond is not buried in the protein core and the solvent accessible surface area (SASA) of the bond increases not satisfying the disulfide bond formation criteria, and it breaks. This allows the possibility that the bonds form in the order specified, allowing the protein fold to its native state (Figure S5D). In the folding pathways, we did not observe any non-native disulfide bond formation, and this shows that protein folding guides disulfide bond formation in agreement with the previous work.^42, 43^

## Conclusions

In summary, using a coarse-grained protein model and molecular dynamics simulations we elucidated the role of disulfide bonds and topological frustration in the folding pathways of lysozyme (Figure 9). The kinetic partitioning^13^ in lysozyme folding pathways to fast and slow routes is due to the two disulfide bonds present in the *α*-domain of the protein. In the slow pathway, a kinetically trapped intermediate state is populated in the late stages of folding due to misalignment in the packing of *α*-helices present in the *α*-domain. The protein has to partially unfold by breaking the inter-helical contacts between the misaligned helices to escape from this trapped state. However, due to the topological restraints imposed by the disulfide bonds in the *α*-domain, the protein is less flexible and it is trapped in the intermediate state for hundreds of milliseconds leading to the slow folding pathway. The schematic describing lysozyme folding pathways is shown in Figure 9, and the energy surface corresponding to this schematic inferred from various experiments was shown by Dobson *et. al.* in Ref.^62^ On disabling the disulfide bonds, the kinetically trapped intermediate state is not populated and the slow folding pathway disappeared. On mimicking lysozyme folding in oxidative conditions by starting the simulations with unfolded conformations without disulfide bonds, we find that it is protein folding, which guides the formation of disulfide bonds.^42, 43^

**Figure 9:**
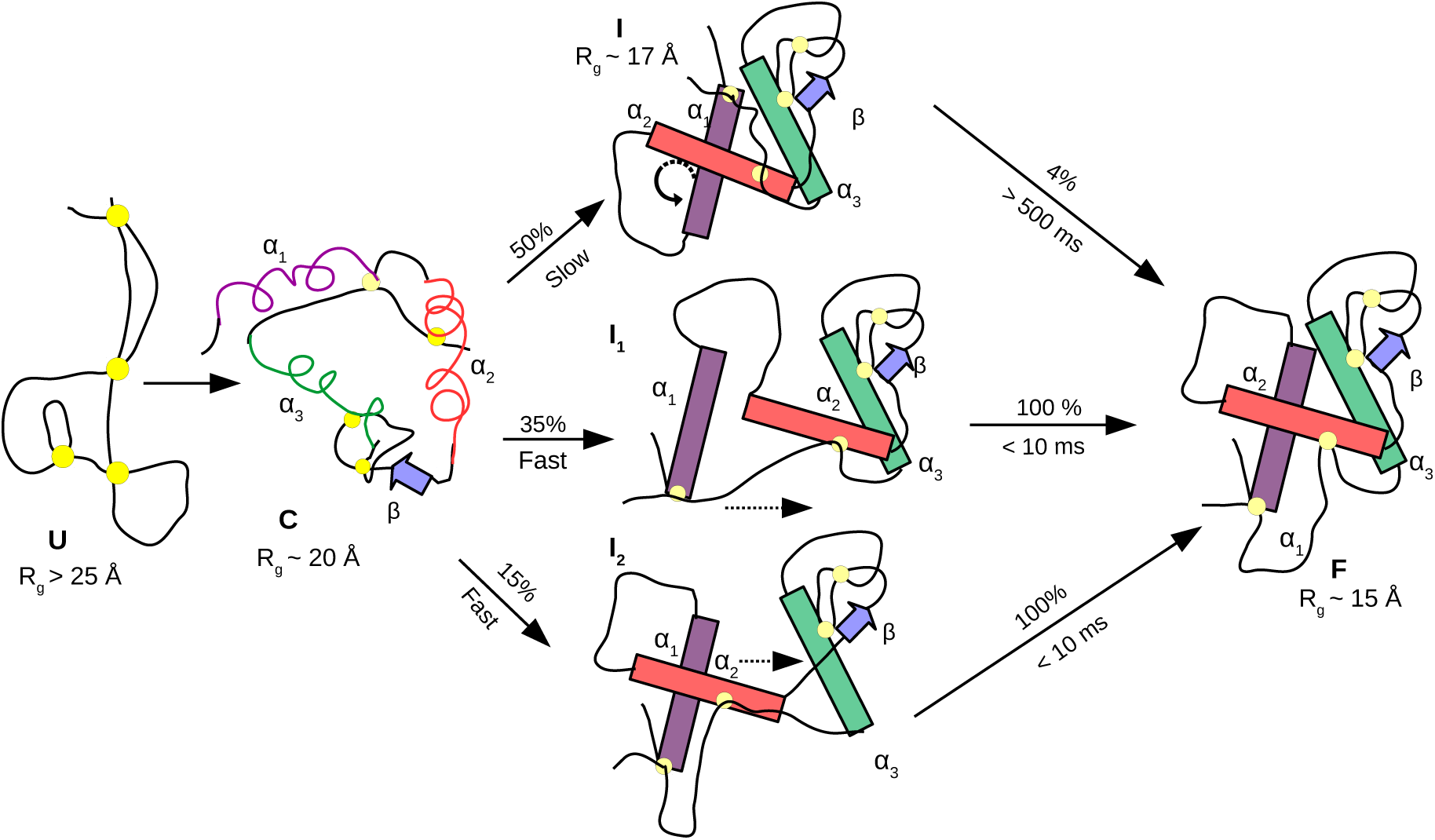
Schematic of dominant folding pathways of hen egg white lysozyme. The three *α*-helices, *α*_1_-*α*_3_, that influence the folding kinetics are shown in purple, red and green, respectively. The triple stranded *β*-sheets, which fold early are shown as one blue arrow. After folding is initiated from an unfolded state (U), the *R_g_* of the protein decreases by ~ 8 Å in size in the first 5 ms due to the folding of *β*-domain and *α*-helices. From this compacted state (C), the folding bifurcates to three different pathways: 1 slow and 2 fast pathways. In the slow pathway, a kinetically trapped state (I) is observed. In this state, the inter-helical contacts *α*_12_ are formed, but the orientation of *α*_1_ relative to *α*_2_ is in the opposite direction compared to the native state. For the protein to fold from this kinetically trapped state, the protein has to backtrack by breaking the *α*_12_ contacts, which will facilitate reorientation of *α*_1_ allowing the protein fold to the native state (F). This rearrangement is restricted by the presence of two disulfide bonds, Cys6-Cys127 and Cys30-Cys115, present in the *α*-domain (Figure 7). In the fast folding pathways, transient intermediate I_1_ or I_2_ are populated. In I_1_, *α*_23_ contacts are formed initially, followed by *α*_12_, *α*_13_ and inter-domain contacts. In I_2_, *α*_12_ contacts are formed initially and helices (*α*_1_ and *α*_2_) have the correct orientation as observed in the native state. In the later stages, *α*_13_, *α*_23_ and inter-domain contacts form and the protein folds to the native state (F).

The kinetically trapped intermediate state populated in the slow folding pathway, which has a lifetime of hundreds of milliseconds, can have implications in the aggregation of lysozyme. In this state, the interface of the *α* and *β*-domains are not completely formed. As the concentration of the kinetically trapped misfolded intermediate increases, it can lead to aggregation through the exposed *β*-sheets.^32, 33^ Experiments^33^ show that mutants, which enhance the population of nonnative states in lysozyme tend to increase the propensity to form fibrils. This shows that disulfide bonds can enhance frustration in the folding landscape of proteins through topological constraints, which can lead to the population of long lived intermediates that have a propensity to aggregate.

## Materials and Methods

To study the folding kinetics of lysozyme we used the SOP-SC (self-organized polymer-side chain)^45, 46^ coarse-grained model for proteins. SOP-SC is a native-centric model ^53^ in which each amino acid in protein is represented by two beads, one for the backbone and the other for side chain. We used the crystal structure^18^ of lysozyme with PDB ID: 1AKI to construct the coarse-grained structures. A detailed description of the SOP-SC model and force field is described in the supplementary information (SI). Brownian dynamics simulations^63^ are performed to probe lysozyme folding kinetics. To mimic the folding of protein in oxidative conditions, where the disulfide bonds form as the protein folds we used the model developed by Qin *et al.*^43^ Details about the simulation methodology and disulfide bond model are described in the SI.

## Acknowledgement

A part of this work is funded by the grants to G.R. and P.C.S. from Science and Engineering Research Board, Department of Science and Technology, India through the grants EMR/2016/001356 and EMR/2015/001605, respectively. The computations are performed using the HPC at TUE-CMS in IISc funded by Nano mission. A.N.M. acknowledges research fellowship from Council of Scientific and Industrial Research, India.

## Supporting Information Available

Detailed description of the protein model, criteria for disulfide bond formation and simulation methods; Figures S1-S6

## Graphical TOC Entry

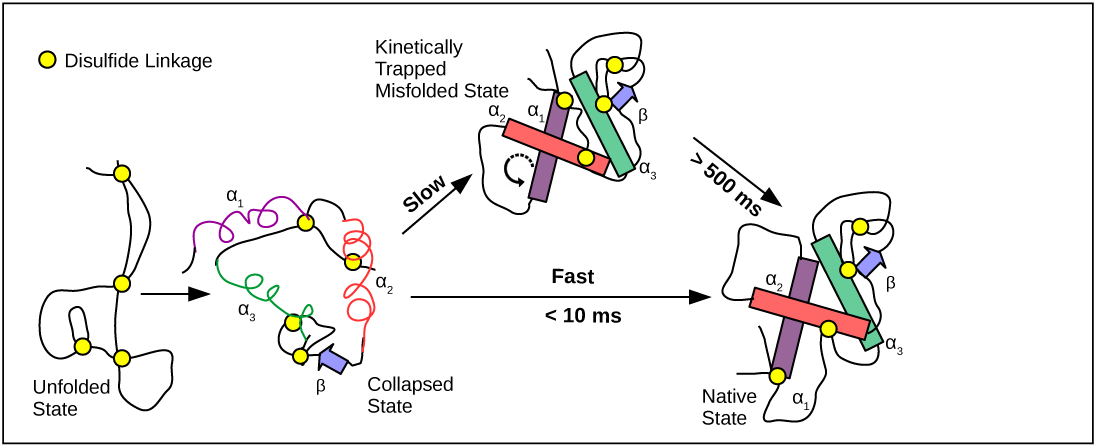

