## Supplementary Information for "Role of Disulfide Bonds and Topological Frustration in the Kinetic Partitioning of Lysozyme Folding Pathways"

### Methods

**Self Organized Polymer-Side Chain (SOP-SC) Model for Proteins:** We have used the native-centric self organized polymer with side chains (SOP-SC) to model lysozyme folding.[1,2] In the SOP-SC model, an amino acid is coarse-grained into two beads representing the back bone and side chain atoms. The centers of the back bone and side chain beads are placed at the $C_\alpha$ position and center of mass of the side chain atoms, respectively. The SOP-SC model for lysozyme is constructed using the crystal structure[3] with PDB ID:1AKI (Figure 1). Hydrogen atoms are added to the protein PDB structure using the program visual molecular dynamics (VMD)[4] before the center of mass of the side chains are calculated.

The Hamiltonian for the SOP-SC model consists of bonded ($E_B$) and non-bonded ($E_{NB}$) terms. $E_{NB}$ is a sum of native ($E_{NB}^N$) and non-native ($E_{NB}^{NN}$) interactions between the beads. Two neighboring beads interact via the bonded potential, $E_B$. If any two beads are separated by more than 2 bonds and if the distance between their centers is less than a cut off distance ($R_c$) in the SOP-SC PDB structure, then the interaction between that pair of beads is native else it is non-native. Native interactions between two consecutive side chain beads are neglected because of their close proximity in both the native and unfolded states. The total energy of a protein conformation in the SOP-SC model described by the set of backbone and sidechain coordinates given by,

$$E_{CG}(\{\mathbf{r}\}) = E_B + E_{NB}^N + E_{NB}^{NN} \tag{S1}$$

The bonds between a pair of beads is modeled using the finite extensible nonlinear elastic (FENE) potential given by

$$E_B = -\sum_{i=1}^{N_B} \frac{k}{2} R_0^2 log\Big(1 - \frac{(r_i - r_{cry,i})^2}{R_0^2}\Big), \tag{S2}$$

where $N_B$ is the total number of bonds in the SOP-SC model of protein, $r_i$ is the distance between the $i^{th}$ pair of beads and $r_{cry,i}$ is the corresponding distance in the native state computed from the SOP-SC PDB structure. The values of $k$ and $R_0$ are listed in Table S1.

The native interactions in the SOP-SC protein model, $E_{NB}^N$, are modeled using a Lennard-Jones type of potential given by

$$\begin{aligned} E_{NB}^N = & \sum_{i=1}^{N_N^{bb}} \epsilon_h^{bb} \Big[\Big(\frac{r_{cry,i}}{r_i}\Big)^{12} - 2\Big(\frac{r_{cry,i}}{r_i}\Big)^{6}\Big] + \sum_{i=1}^{N_N^{bs}} \epsilon_h^{bs} \Big[\Big(\frac{r_{cry,i}}{r_i}\Big)^{12} - 2\Big(\frac{r_{cry,i}}{r_i}\Big)^{6}\Big] \\ & + \sum_{i=1}^{N_N^{ss}} 0.5 \times 300 k_B \times (0.7 - \epsilon_i^{ss}) \Big[\Big(\frac{r_{cry,i}}{r_i}\Big)^{12} - 2\Big(\frac{r_{cry,i}}{r_i}\Big)^{6}\Big], \end{aligned} \tag{S3}$$

where $N_N^{bb}$, $N_N^{bs}$ and $N_N^{ss}$ denote the number of native contact pairs between backbone -

backbone, backbone - side chain, and side chain - side chain, respectively. $r_i$ is the distance between $i^{th}$ pair of beads and $r_{cry,i}$ is the corresponding distance in the SOP-SC crystal structure. $\epsilon_h^{bb}$, $\epsilon_h^{bs}$ and $\epsilon_i^{ss}$ denote the strength of backbone - backbone, backbone - side chain, and side chain - side chain interactions, respectively. The values of $\epsilon_i^{ss}$ are taken from the Betancourt-Thirumalai statistical potential,[5] $\epsilon_h^{bb}$ and $\epsilon_h^{bs}$ are listed in Table S1, and $k_B$ is the Boltzmann constant.

The non-native interactions, $E_{NB}^{NN}$, are purely repulsive interactions and are given by

$$E_{NB}^{NN} = \sum_{i=1}^{N_{NN}} \epsilon_l \Big(\frac{\sigma_i}{r_i}\Big)^6 + \sum_{i=1}^{N_{ang}^{bb}} \epsilon_l \Big(\frac{\sigma^{bb}}{r_i}\Big)^6 + \sum_{i=1}^{N_{ang}^{bs}} \epsilon_l \Big(\frac{\sigma_i^{bs}}{r_i}\Big)^6, \tag{S4}$$

where $N_{NN}$ is the total number of non-native pairs, $\sigma_i$ is sum of the radii of $i^{th}$ pair of beads, and $\sigma^{bb}$ is the diameter of backbone bead. The second and third terms on the right hand side of eq. S4 mimic the bond angle potential between beads. $N_{ang}^{bb}$ is the number of bond angles between backbone beads, and $N_{ang}^{bs}$ is the number of bond angles involving backbone and side chain beads. $\sigma_i^{bs} = f[\sigma^{bb} + \sigma_i^{sc}]/2.0$, where $\sigma_i^{sc}$ is the diameter of the side chain bead corresponding to the $i^{th}$ bond angle involving a backbone and side chain bead. $f(= 0.8)$ is a rescaling factor used to minimize the repulsion between the beads. Values of the side chain radii are given in Table S3.[6,7]

**Simulations:** To provide a realistic description of the folding kinetics we performed Brownian dynamics simulations and the equations of motion are integrated using the Ermak-McCammon algorithm,[8] $\vec{r_i}(t + h) = \vec{r_i}(t) + \frac{h}{\zeta}\vec{F_c} + \vec{\Gamma}$. Here $\vec{\Gamma}$ is a random force with a Gaussian distribution with mean zero and variance $\langle \Gamma(h)^2 \rangle = \frac{2k_BTh}{\zeta}$. The friction coefficient $\zeta = 50\ m/\tau_H$ approximately corresponds to the value in water and $h = 0.005\ \tau_H$. In the simulations, the characteristic unit of length $a = 1$ Å, energy $\epsilon = 1\ kcal/mole$, and mass $m = 1.8 \times 10^{-22}\ g$ (typical mass of the bead). In Brownian dynamics, simulation time is mapped into real time using $\tau_H \approx \frac{\zeta_H a^2}{k_BT} = \frac{(\zeta_H \tau_L/m)\epsilon}{k_BT}\tau_L \approx 43\ ps$.

**Simulations Mimicking Oxidative Folding of Lysozyme:** To understand the order of formation and rupture of disulfide bonds in a buffer containing the reducing agent glutathione, we adopted the method developed by Qin *et al.*[9] to two bead model. In this model formation or rupture of disulfide bonds are determined by three criteria - (i) proximity of Cys side chains ($r$) (ii) relative orientation of Cys residues ($\theta_1$ and $\theta_2$) and (iii) exposure of disulfide bonds - quantified by number of residues within a radius $R_s = 7$ Å from the center of the disulfide bonds ($n$) (Figure S6). These values are calculated at each step of the simulation ($x^i$) and are compared to the values in the native state ($x^{nat} \pm \delta_x$). The native state values are calculated from equilibrium trajectories at a temperature where the folded state is stable (T = 320 K). A disulfide bond between a pair of Cys residues is allowed to form in a protein conformation, if the following criteria is satisfied: $r^i < r^{nat} + \delta_r$, $\theta^i < \theta^{nat} + \delta_\theta$ and $n^i > n^{nat} - \beta\delta_n$. In the present work, we used $\beta = 1.0$ for oxidation and $\beta = 1.5$ for reduction. Once the criteria for the disulfide bond formation is satisfied for a pair of Cys residues, a FENE potential (eq. S2) is added between the side chain beads of those residues to mimic the disulfide bond.

Table S1: Energy function parameters for the SOP-SC model

| Parameters | value |
|---|---|
| $R_o$ | 2.25 Å |
| $k$ | 20 kcal/(mol. Å$^2$) |
| $R_c$ | 8 Å |
| $\epsilon_h^{bb}$ | 0.38 kcal/mol |
| $\epsilon_h^{bs}$ | 0.38 kcal/mol |
| $\epsilon_l$ | 1.0 kcal/mol |
| $\sigma^{bb}$ | 3.8 Å |

Table S2: Protein SOP-SC model Parameters

| Parameters | Without Disulfide | With Disulfide |
|---|---|---|
| $N_B$ | 257 | 261 |
| $N_N^{bb}$ | 377 | 373 |
| $N_N^{bs}$ | 988 | 980 |
| $N_N^{ss}$ | 437 | 433 |
| $N_{NN}$ | 30583 | 30583 |
| $N_{ang}^{bb}$ | 127 | 127 |
| $N_{ang}^{bs}$ | 256 | 264 |

Table S3: Side chain and backbone radii of amino acids based on partial molar volumes

| Residue | Radius (Å) |
|---|---|
| *Gly* | 0 |
| *Ala* | 2.52 |
| *Val* | 2.93 |
| *Leu* | 3.09 |
| *Ile* | 3.09 |
| *Met* | 3.09 |
| *Phe* | 3.18 |
| *Pro* | 2.78 |
| *Ser* | 2.59 |
| *Thr* | 2.81 |
| *Asn* | 2.84 |
| *Gln* | 3.01 |
| *Tyr* | 3.23 |
| *Trp* | 3.39 |
| *Asp* | 2.79 |
| *Glu* | 2.96 |
| *His* | 3.04 |
| *Lys* | 3.18 |
| *Arg* | 3.28 |
| *Cys* | 2.74 |
| *backbone* | 1.9 |

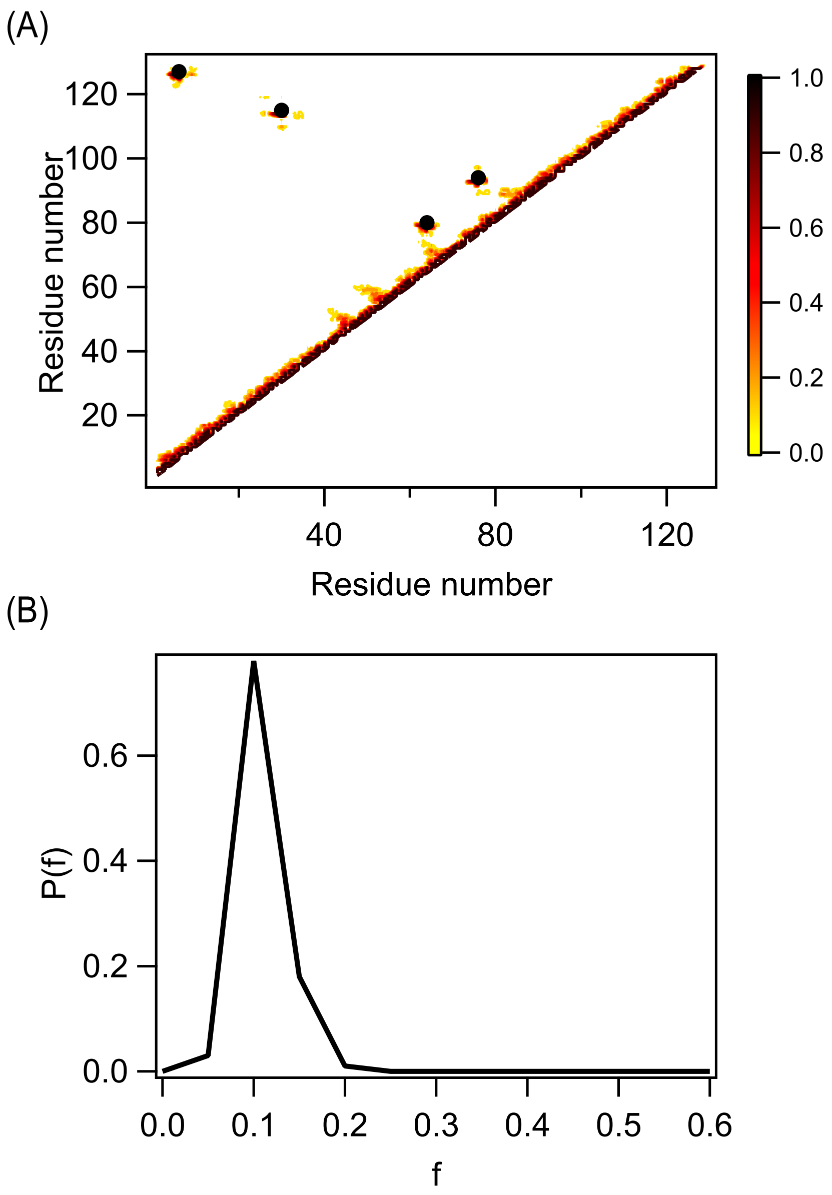


Figure S1: (A) Contact map of lysozyme unfolded ensemble obtained at T = 420 K. Protein conformations are used from this ensemble to start the Brownian dynamics folding simulations. (B) Probability distribution of the fraction of native contacts in the 100 initial unfolded conformations used to initiate folding simulations. Both the contact map and distribution of native contacts show that protein conformations in the unfolded ensemble lack native structure.

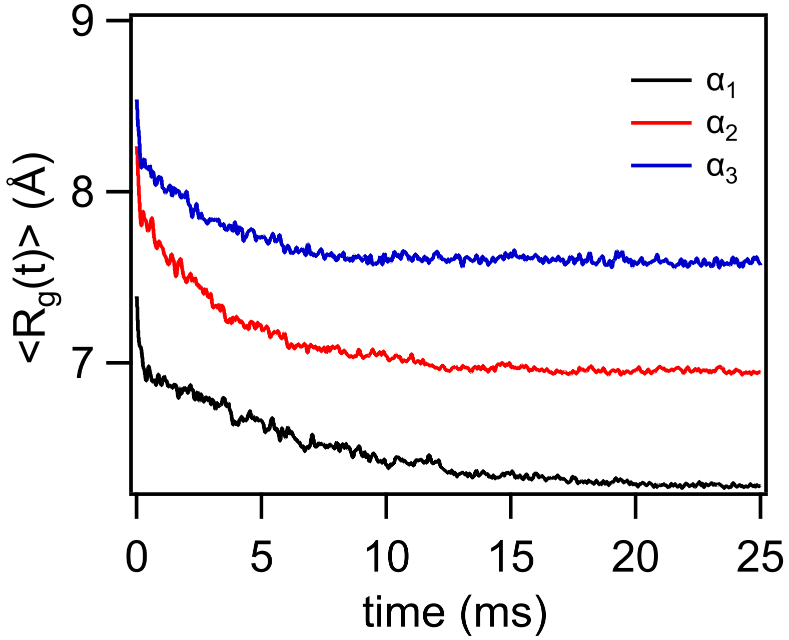


Figure S2: Time evolution of the ensemble average of radius of gyration, $\langle R_g(t) \rangle$, of $\alpha$-helices. Within the collapse time ($\sim$ 5 ms), $\langle R_g(t) \rangle$ of the helices is close to the native state $R_g$. This implies that the formation of helices contributes to the reduction in lysozyme size during collapse.

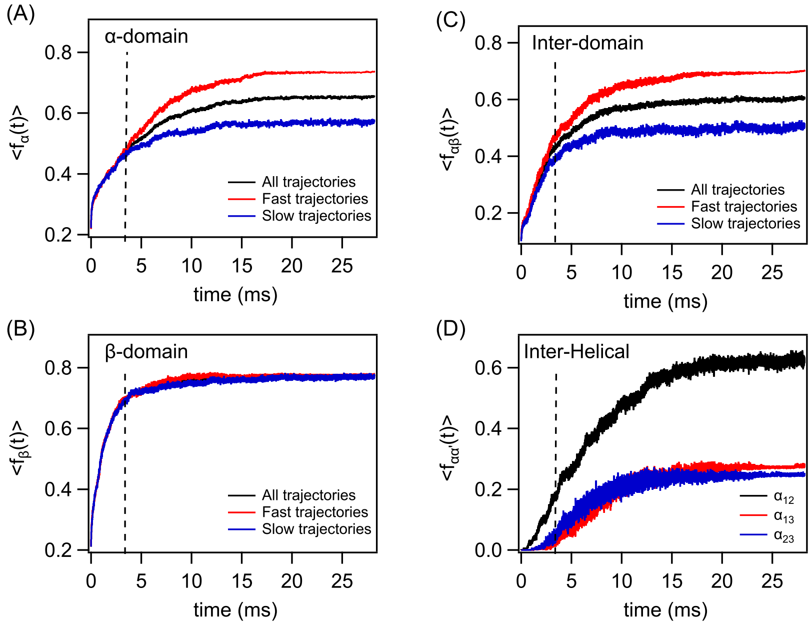


Figure S3: Time evolution of the ensemble average of the fraction of native contacts in the $\alpha$-domain (A), $\beta$-domain (B), interface of $\alpha\beta$ domain (C), and between helices, $\alpha_1 - \alpha_3$ (D) in fast and slow folding pathways. The dotted line refers to the time scale for collapse in lysozyme. During the collapse, $\beta$-domain is completely formed, and a significant fraction of the inter-domain contacts are formed. It is mostly the inter-helical contacts that are not formed.

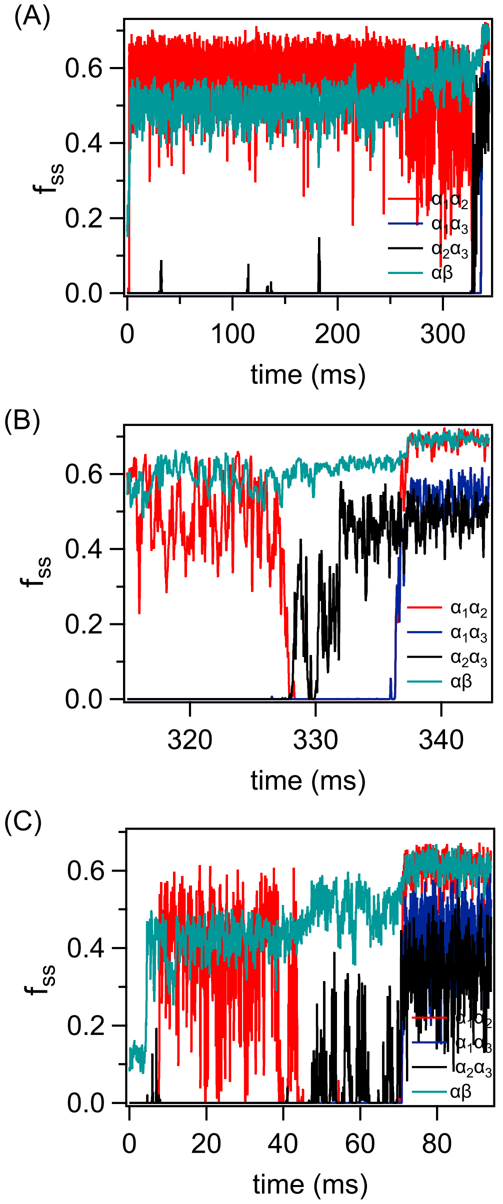


Figure S4: (A) Lysozyme folding using the slow pathway at T = 320 K. The inter-helical and inter-domain contacts are shown as a function of time. For the protein to fold from the kinetically trapped intermediate state, $\alpha_1\alpha_2$ helical contacts between the misaligned helices should breakdown, allowing the helices to realign to the correct orientation. (B) The last 50 ms of the trajectory, which shows the breakdown in $\alpha_1\alpha_2$ contacts followed by the reformation of all the helical contacts in the correct orientation. (C) Lysozyme folding using the slow pathway at T = 380 K. Similar folding mechanism is observed.

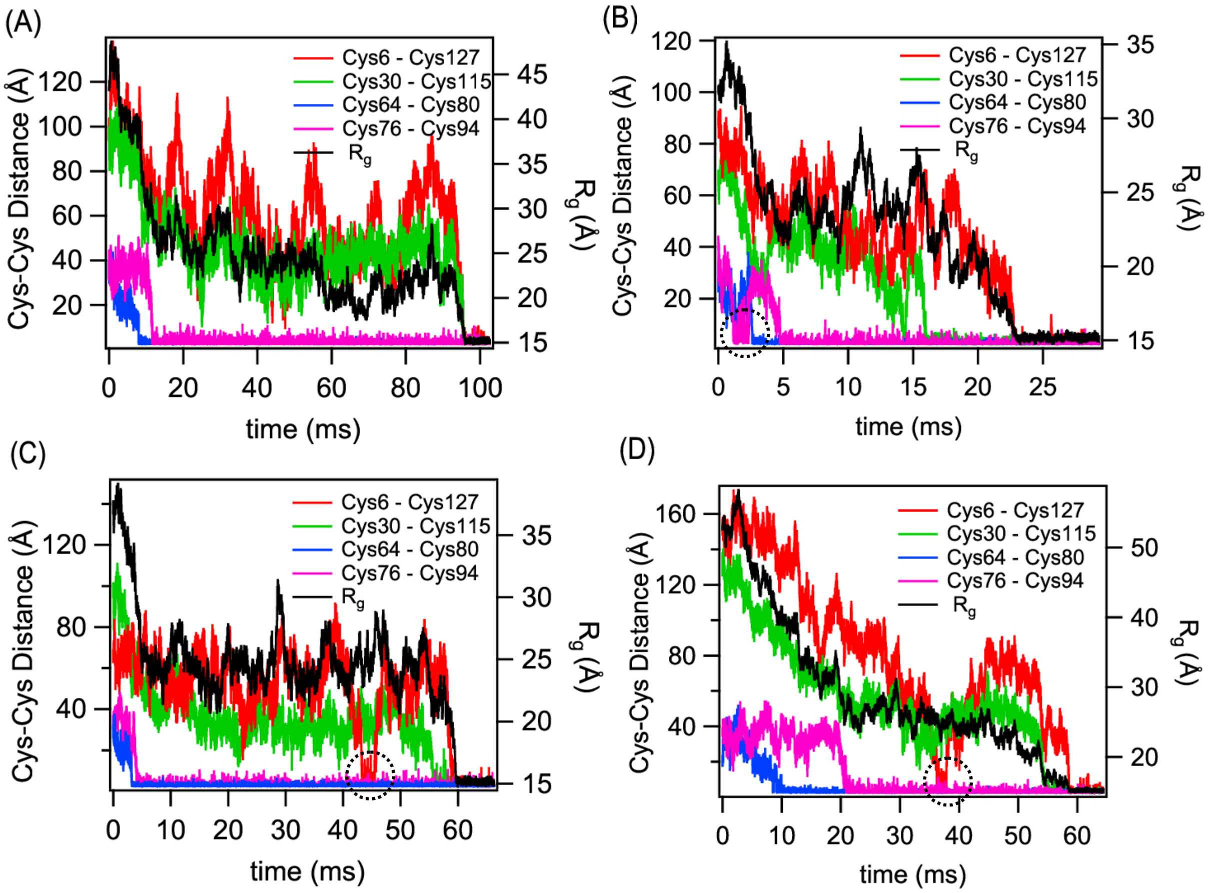


Figure S5: Lysozyme folding in oxidative conditions. (A) Folding trajectory where Cys6 - Cys127 and Cys30 - Cys115 disulfide bonds present in the $\alpha$-domain are simultaneously formed. This is observed in 6 out of the 100 folding trajectories. (B,C,D) Folding trajectories where the sequence of disulfide bond formation, Cys64-Cys80 $\rightarrow$ Cys76-Cys94 $\rightarrow$ Cys30-Cys115 $\rightarrow$ Cys6-Cys127 is not followed. The disulfide bond, which forms ahead of its precursor in the sequence breaks down to follow the sequence. The breaking down of the bond is highlighted in circles.

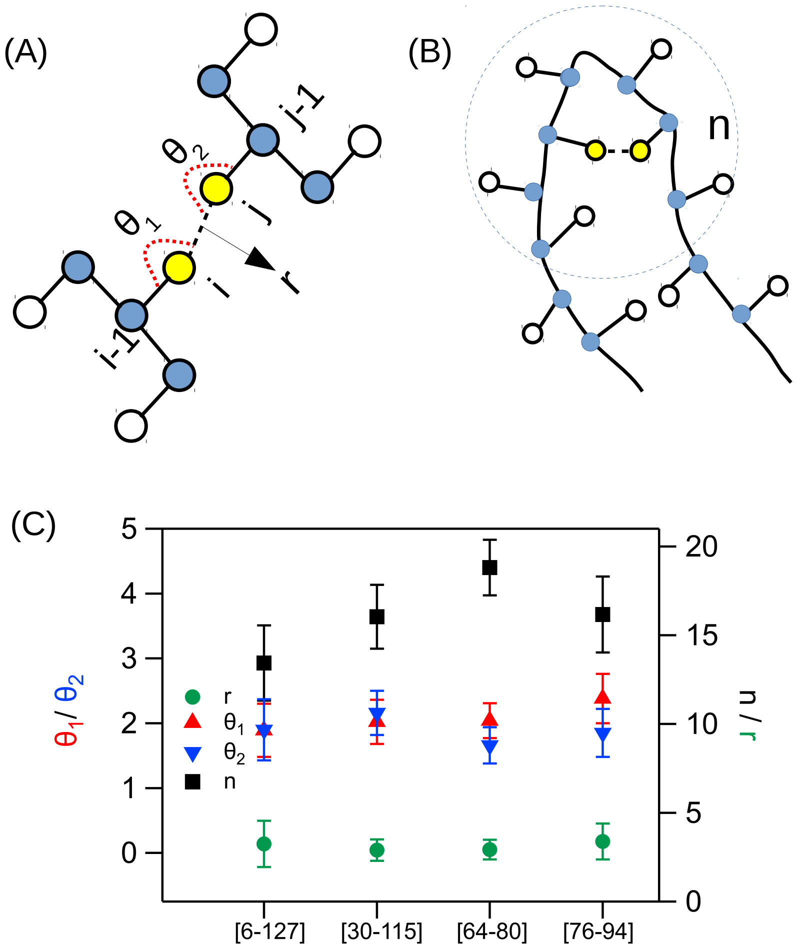


Figure S6: Criteria for disulfide bond formation. (A) The distance, $r$, between two Cys residues $i$ and $j$, and the angles, $\theta_1$ and $\theta_2$, the bond makes with the Cys backbone beads are used to define the disulfide bond. If the values of $r$, $\theta_1$ and $\theta_2$ are within the fluctuations of its native state values, then a disulfide bond is allowed to form between $i$ and $j$. (B) The fourth criteria that has to be satisfied for disulfide bond formation is based on the exposure of disulfide bonds. If the number of residues, $n$, within a cutoff distance, $R_s = 7.0$ Å, around the disulfide bonds (dotted circle) is less than the native state values, then the disulfide bond is considered to be exposed to the reducing agents. (C) The average values of the parameters obtained from equilibrium simulations at T$=$ 320 K with standard deviations are shown for the four disulfide bonds.
